## Supplementary figures and images for "Characterization of RNA interference in the model cnidarian *Nematostella vectensis* reveals partial target silencing but lack of small RNA amplification"

### Supplementary Figure 1 .tif

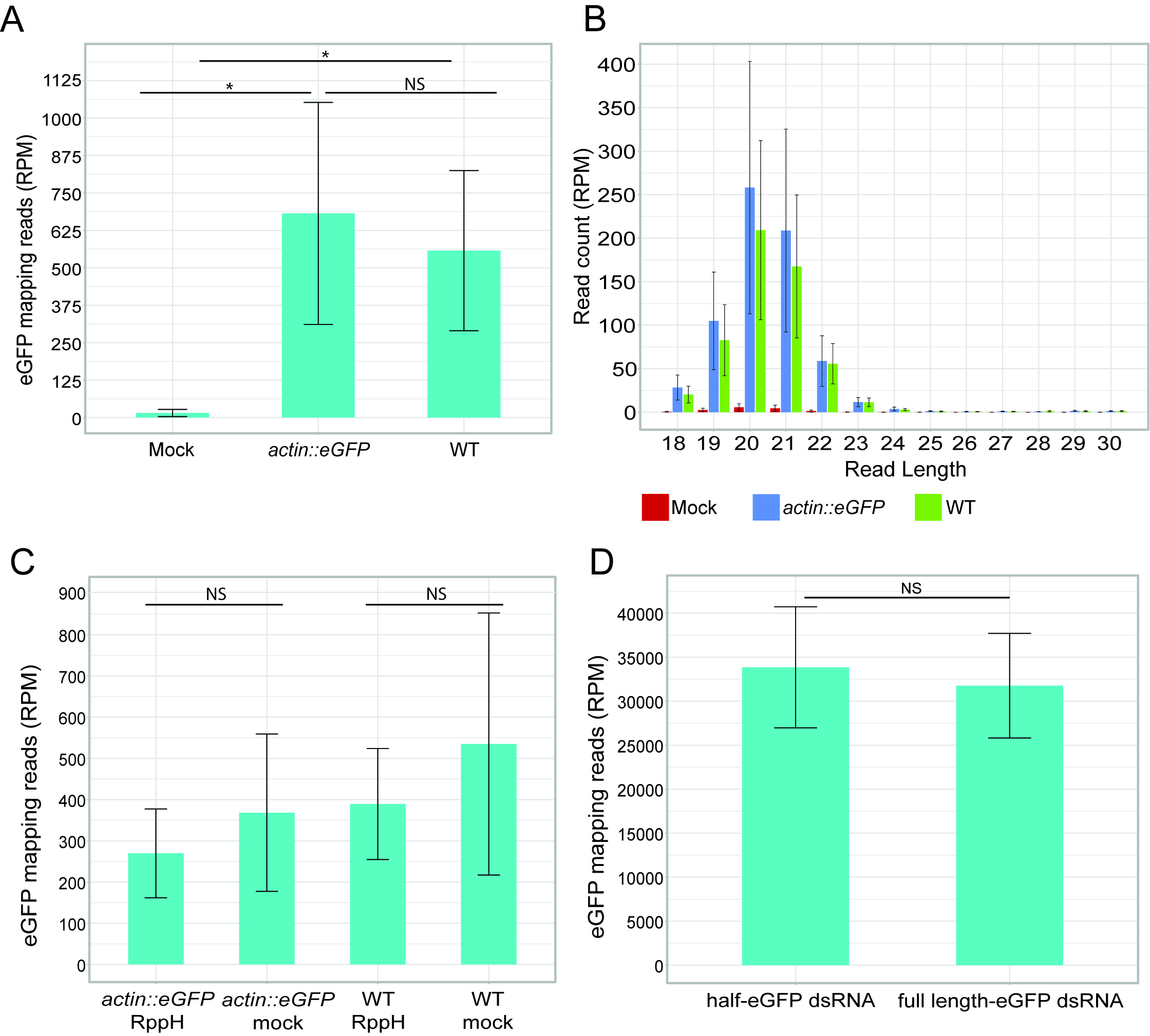

### Supplementary Figure 2.tif

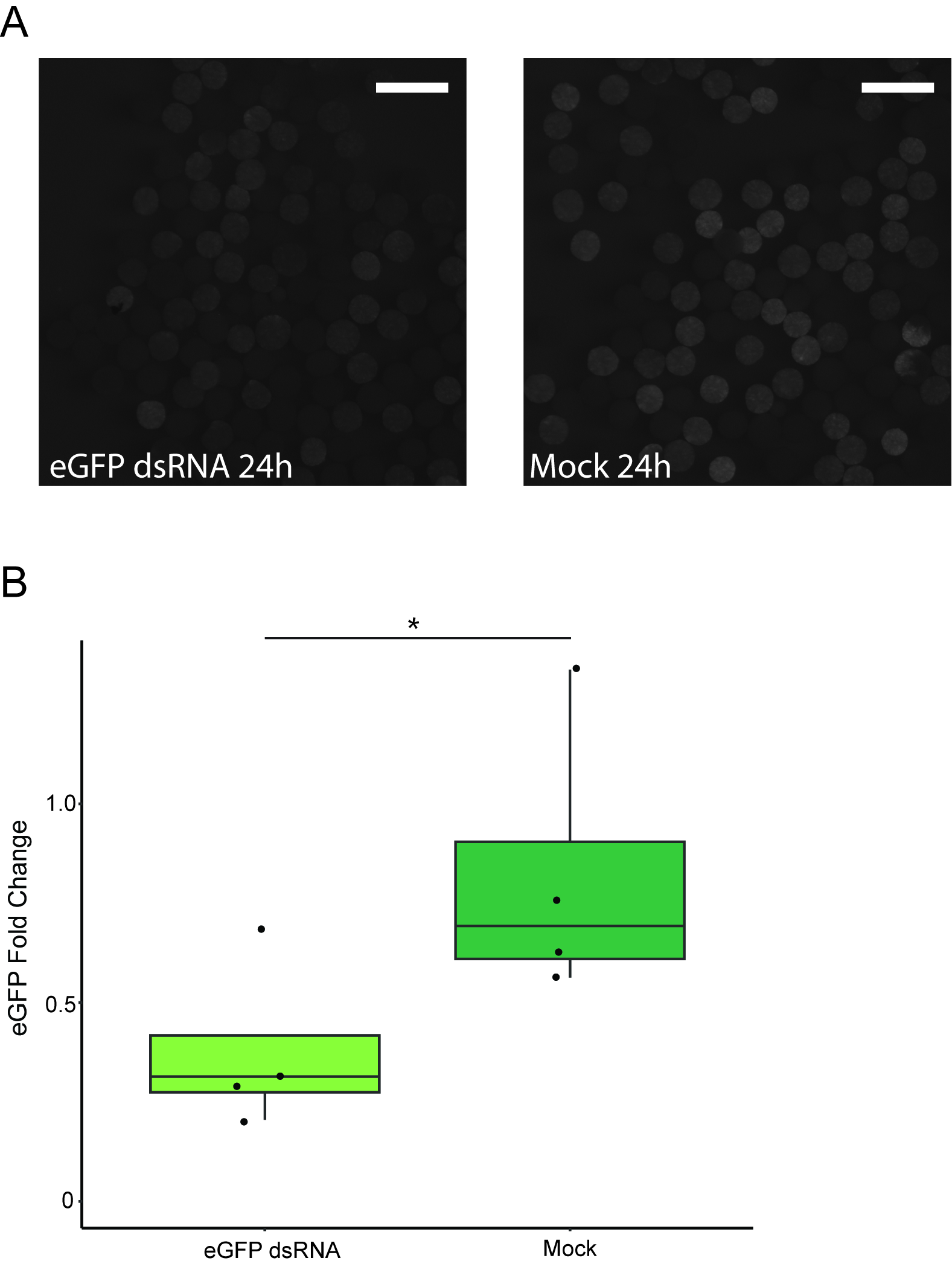

### Supplementary Figure 3.tif

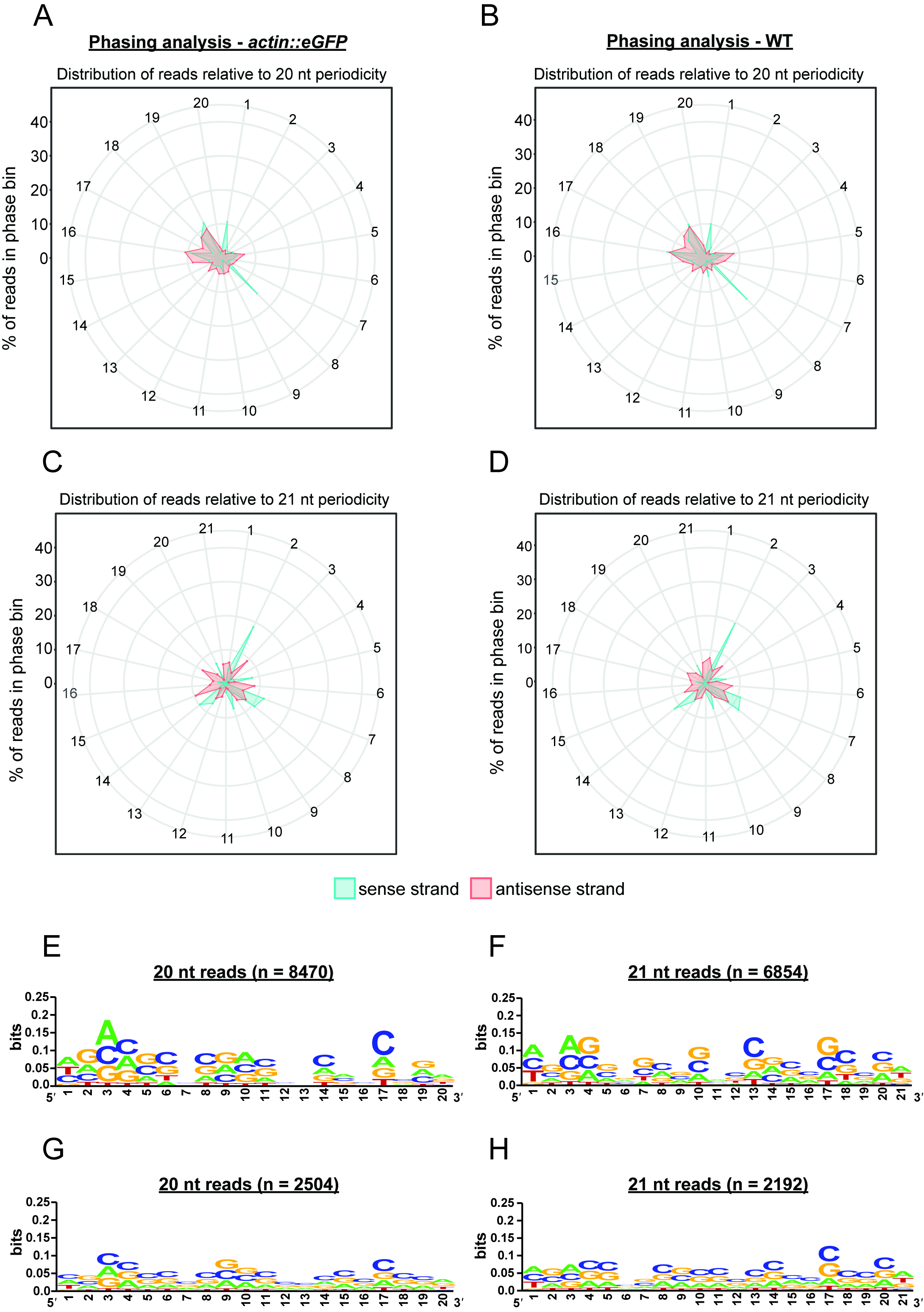

### Supplementary Figure 4.tif

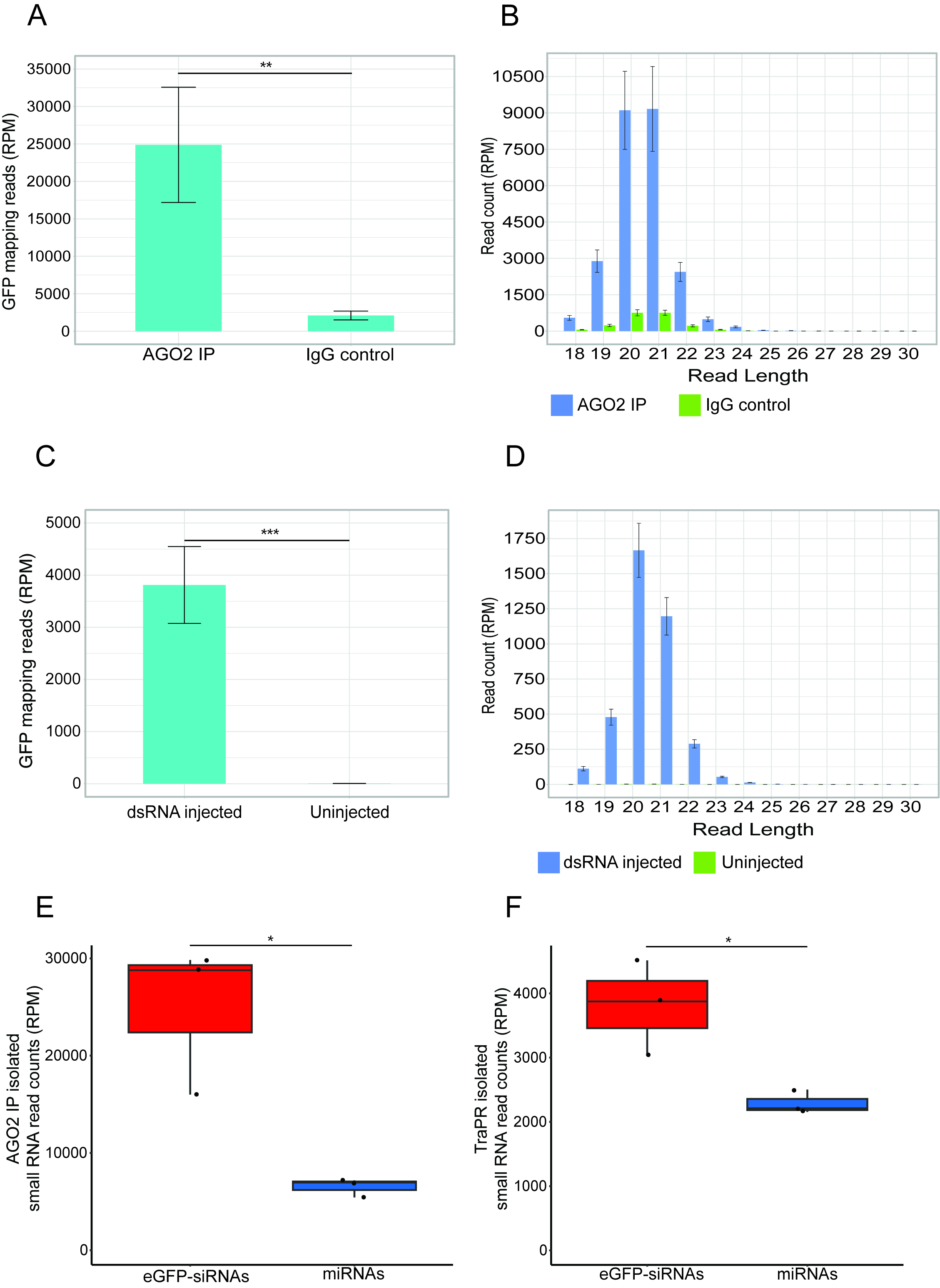

### Supplementary Figure 5.tif

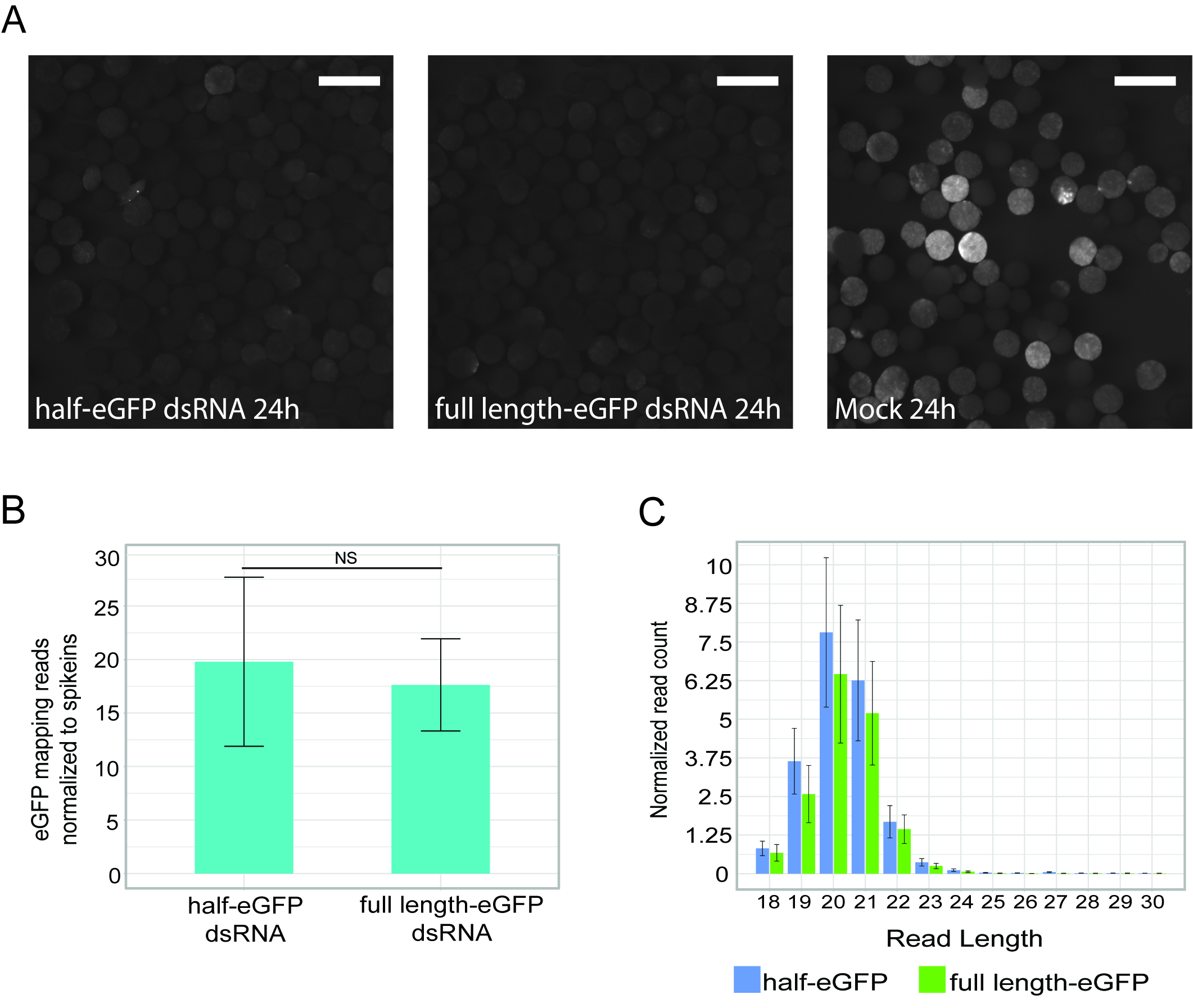

### Supplementary Figure 6.tif

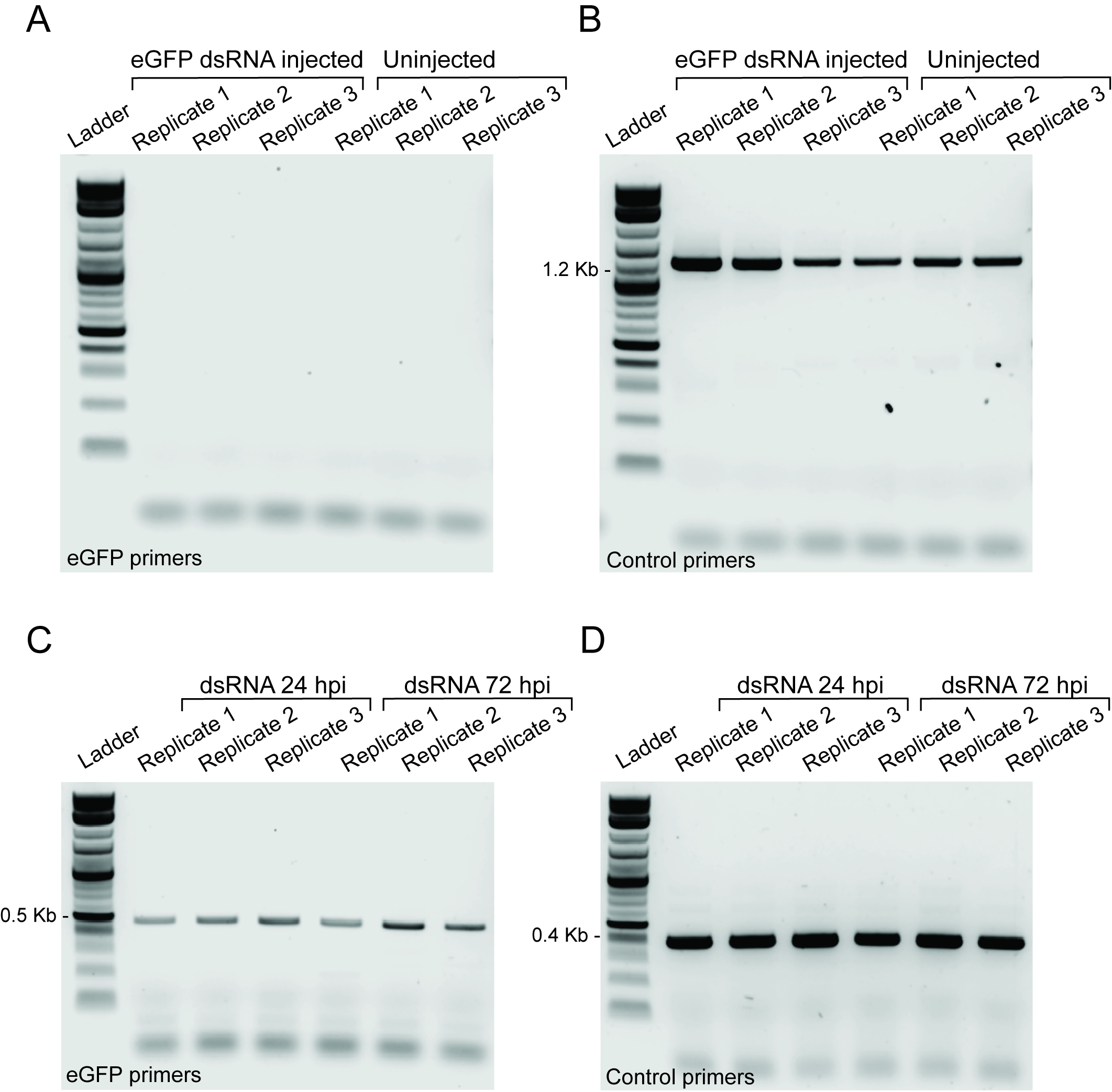

### Supplementary Figure 7.tif

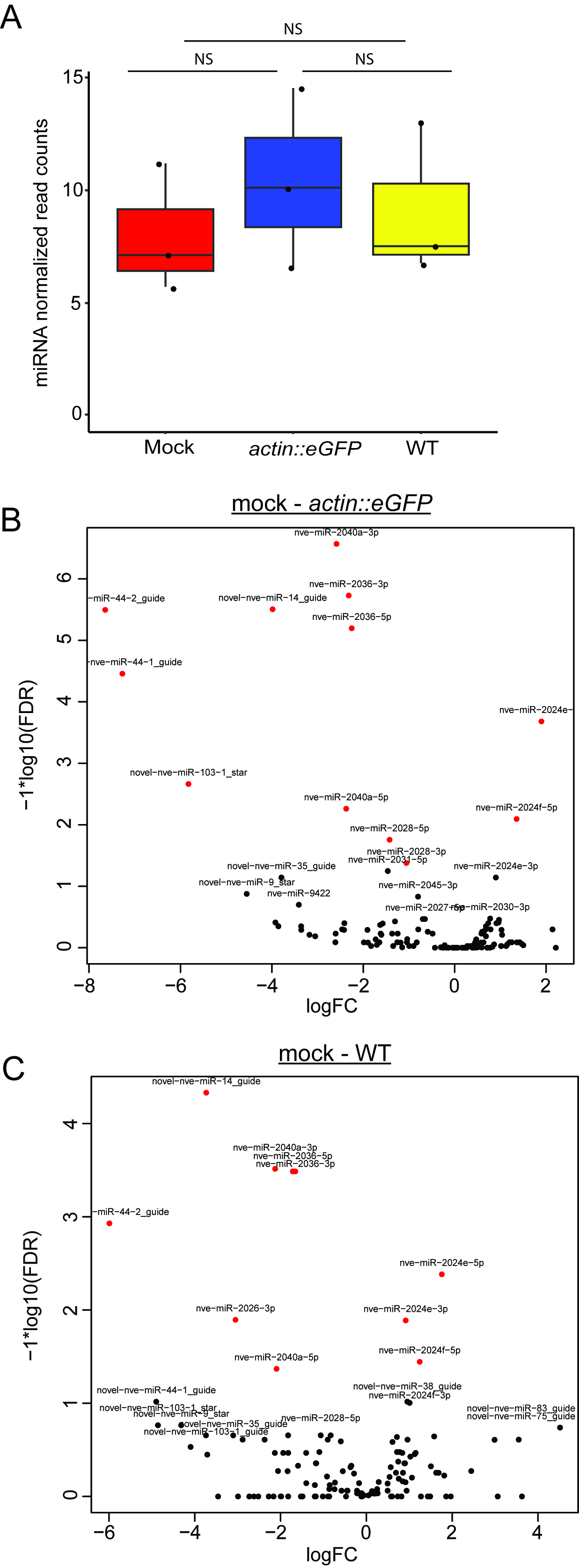

### Supplementary Figure 8.tif

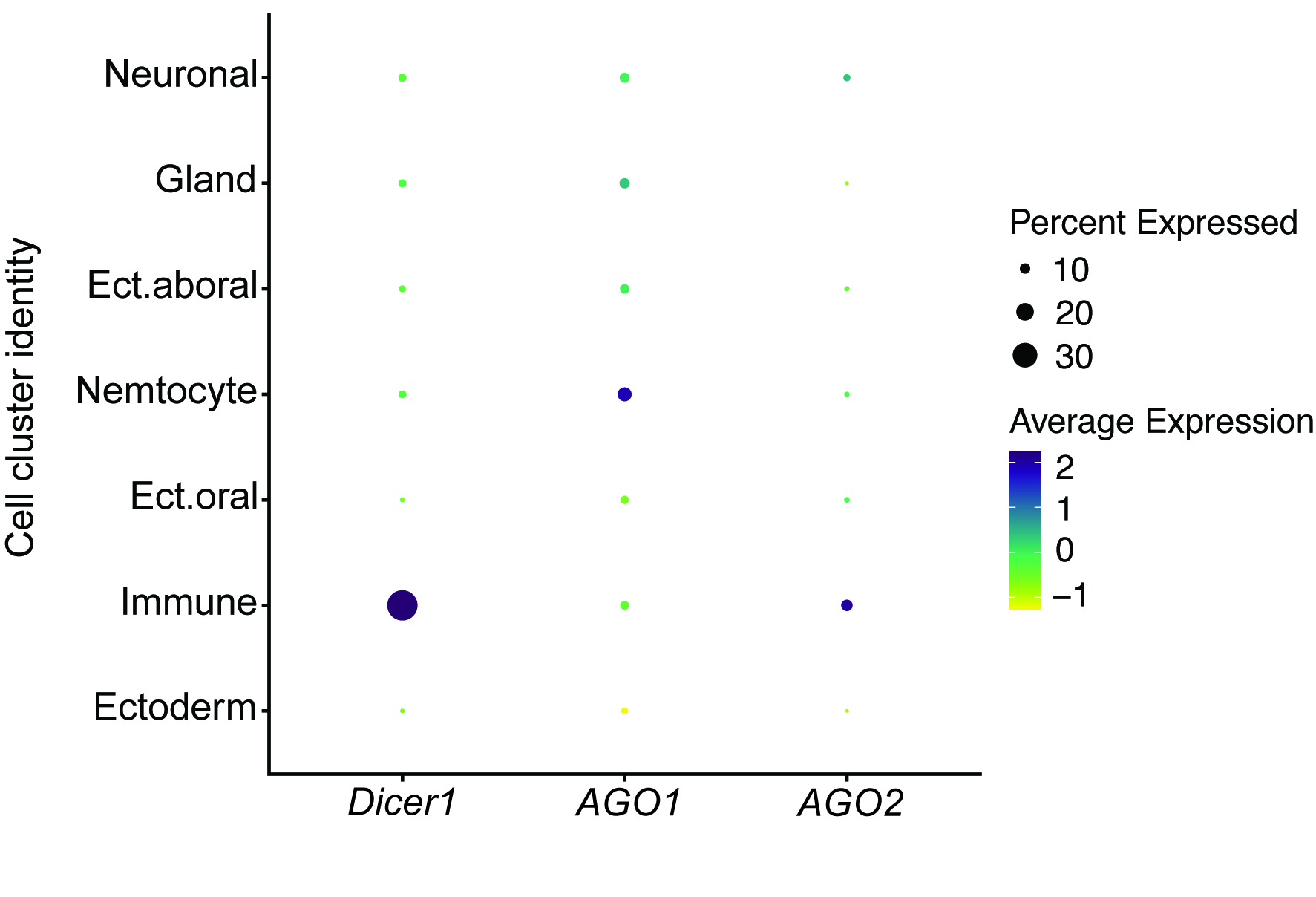
